# Effects of binge-like ethanol drinking on nest building behavior in mice

**DOI:** 10.1101/2025.06.03.657694

**Authors:** André Lucas Silva-Borges, Amanda M. Barkley-Levenson

## Abstract

Nest building is a natural behavior that can readily be analyzed in mice in the home cage environment. Nest building is involved in thermoregulation, positive motivational states, and motor function, and alterations in this behavior have been proposed as an index for ethanol withdrawal severity in mice. However, nest building outcomes after voluntary ethanol consumption have not been examined. Here, we tested male and female C57BL/6J mice on a 4-day drinking in the dark (DID) paradigm of binge-like drinking with either ethanol or a water control and analyzed nest scores at two timepoints (48 hours and 7 days) after the last DID session. At 48 hours after the last DID session, there were no differences between the two groups in nest quality. At 7 days after DID, ethanol-drinking animals showed significantly lower nest scores than the water group (z = -2.369, *p*=0.030). No differences were found between the ethanol- and water-drinking groups in locomotor activity or anxiety-like behavior at this timepoint in an open field test, indicating that nest building deficits in the ethanol group were likely not due to underlying differences in these behaviors. Together, these results validate the use of nest building as a naturalistic assessment of post-ethanol behavioral changes following voluntary binge-like drinking.

## Introduction

Binge drinking is defined as a pattern of alcohol (ethanol) drinking that results in blood ethanol concentrations (BEC) of 0.08% or higher, which typically equates to five or more drinks on one occasion for men and four or more for women. In the United States, roughly 28.7% of young adults report weekly binge drinking, and 90% of adults who drink excessively report binge drinking^1^. This drinking pattern can lead to significant individual and societal costs and represents a serious public health concern. Additionally, binge drinking is associated with a higher risk of developing an alcohol use disorder and related problems such as withdrawal symptoms^2^. Ethanol withdrawal symptoms can include negative affective changes such as anxiety, depression, and agitation^3^.

In mice, previous studies have shown that previous exposure to ethanol injection or ethanol vapor inhalation can cause lasting changes in affective and goal-directed behaviors, including nest building behavior^4^. Nest building is a highly conserved behavior in mice that is seen in both the wild and in laboratory animals^5^. Nest building is relevant for thermoregulation, predator defense, and parental care, and has been proposed to represent a potential model of an “activity of daily living” in mice^6,7^. Both male and female mice build nests and sex differences are not consistently reported, though this may vary depending on age and genetic background^,8,9,11^. Nest building is a sensitive index of general well-being in mice and has been shown to be impacted by physiological states such as stress and sickness, as well as specific brain lesions and genetic mutations^5,10^. Several studies have also reported nest building deficits during ethanol withdrawal, following either acute ethanol injection or ethanol vapor inhalation exposure^4,11^. Nest building may therefore represent a simple, non-invasive means of assessing ethanol withdrawal severity and post-ethanol behavioral changes in mice.

In the present study, we sought to build upon these previous findings to determine whether voluntary binge-like ethanol drinking is sufficient to produce lasting deficits in nest building. Mice were tested on a 4-day drinking in the dark (DID) paradigm with either 20% ethanol or water, and nest quality was evaluated at 48 hours (early post-drinking period) and 7 days (late post-drinking period) after the final binge-like drinking day. We hypothesized that this paradigm would promote the same deficits previously seen in nest building behavior after other types of ethanol exposure, therefore validating the use of this test for the evaluation of abstinence-associated behavioral changes after voluntary ethanol consumption.

## Material and methods

### Animals and husbandry

Forty adult male and female C57BL/6J mice (n=10/sex/drinking group) were purchased from the Jackson Laboratory (Bar Harbor, ME, USA) and arrived at our facility at 8 weeks old. Mice were allowed to acclimate to the animal facility and a 12hr/12hr reverse light-dark cycle with lights off at 08:00 for two weeks prior to the start of testing. One week before the start of testing, mice were singly-housed in standard shoebox cages on Sani-chip bedding with a sipper tube water bottle. They were fed 5001 LabDiet and had access to both food and water *ad libitum*.

Mice were also provided with a single 2” square cotton nestlet as enrichment to allow familiarization with the nest building material. All experimental procedures were approved by the Institutional Animal Care and Use Committee at the University of New Mexico Health Sciences Center, were conducted in accordance with the NIH Guidelines for the Care and Use of Laboratory Animals, and are reported here according to ARRIVE guidelines.

### Drinking in the Dark (DID)

Prior to the start of the experiment, mice were pseudorandomly assigned to either the ethanol or water drinking group. On each day of the DID procedure, mice were weighed approximately 30 min before the start of the drinking session. Three hours after lights-off, water bottles were removed and replaced by 10 ml tubes fitted with a ball bearing metal sipper and containing either 20% ethanol (v/v in water) or water depending on group assignment. On Days 1-3, fluid levels were recorded, and tubes were left in place for 2 hours, after which fluid levels were recorded again. Tubes were then removed and water bottles were returned to the cages. On Day 4, the procedure was the same as on the previous days, except that drinking tubes were left in place for 4 hours, and fluid levels were recorded after 2 and 4 hours.

### Nest Building Tests

24 hours after the end of the final DID session, the existing nesting material was removed from the cage and replaced with a fresh cotton nestlet. Mice were left undisturbed in the home cage for 24 hours after nesting material replacement, and then the nest quality was scored. Six days after the final DID session, the nesting material was again removed and replaced with a fresh nestlet, and nests were again scored 24 hours after replacement. Nest scoring was based on the scale previously described by Deacon^6^ and used the following criteria: untouched nestlets received a score of 1, shredded nestlets that had been moved and scattered without a centralized nest site received a score of 2, centralized nest sites without any walls received a score of 3, centralized nest sites with walls received a score of 4, and centralized nest sites with fully domed walls received a score of 5. Nests with qualities fitting in multiple scoring categories were awarded half points between the identified categories.

### Open Field Test

The open field test was performed 7 days after the final DID session, immediately following the second nest score assessment. The test apparatus consisted of an arena made of acrylic transparent walls and white floor (27 × 27 × 20cm; Med Associates, St. Albans, VT) and placed inside of an illuminated sound-attenuating cubicle. Infrared light sensors separated by 1.5 cm intervals were used to automatically detect horizontal and vertical activity at 1.5 and 6 cm above the floor level, respectively. The center space was defined as 8×8 beams (12.72 cm × 12.72 cm). On the experiment day, the animals were moved to the experiment room and allowed to acclimate for 1 hour. Mice were then placed individually into the center of the arena and allowed to explore freely for 15 minutes. Locomotor activity (distance traveled in cm) and the time spent in the center of the arena were automatically recorded. Between animals, the chambers were cleaned with 10% isopropyl alcohol to eliminate odor cues.

### Statistical analyses

Statistical analyses were carried out using SPSS (IBM; version 29.0.1.0(171)). Alcohol intake was analyzed by repeated measures analysis of variance (ANOVA) with sex as a factor. Total intake on the last day of DID was analyzed via one-way ANOVA, also including sex as a main factor. Mann-Whitney U tests were used to assess drinking group and sex effects on nest scores at 48 hours and 7 days. Separate analyses were conducted for each timepoint due to our *a priori* hypothesis that nest building during the early post-drinking period (acute “hangover” phase) and the later post-drinking period are impacted by different underlying effects of previous ethanol intake. Total locomotor activity (distance traveled in cm) and percent of total time spent in center of the chamber were also analyzed using two-way ANOVA with factors of treatment group and sex. Data are shown collapsed on sex when no significant treatment group x sex interaction was found. Spearman’s correlation was used to determine if there was a significant relationship between cumulative DID ethanol intake and nest score at either timepoint. Significance was set at α=0.05 for all tests.

## Results

### Drinking in the Dark and Nest Building

Figure 1 shows experimental timeline (Fig. 1A) and average g/kg intake of ethanol on each day of the DID test (Fig. 1B). There was a significant main effect of sex for the ethanol intake repeated measures analysis [*F*(1,18)=7.530, *p*=0.013], with the females drinking more than the males, which was also seen when the total intake on the last day was analyzed [*F*(1,18)=4.526, *p*=0.047]. This is consistent with the existing literature regarding the levels of ethanol intake in C57BL/6J mice^12-15^. Mice in the water drinking group did not show any significant sex differences in ml/kg intake. Figure 2 shows nest scores on each of the post-drinking assessments. At 48 hours after the last drinking session, the treatment groups did not significantly differ in nest score (z = -0.385, *p*=0.708; Fig. 2A). At 7 days following drinking, however, there the groups did significantly differ, with the ethanol-drinking animals having poorer nest quality scores than the water-drinking control animals (z = -2.369, *p*=0.030; Fig. 2B). There were no significant effects of sex on either nest building test, and no significant correlations between cumulative DID ethanol intake and nest scores at either test (*p*≥0.05 for all).

**Figure 1.**
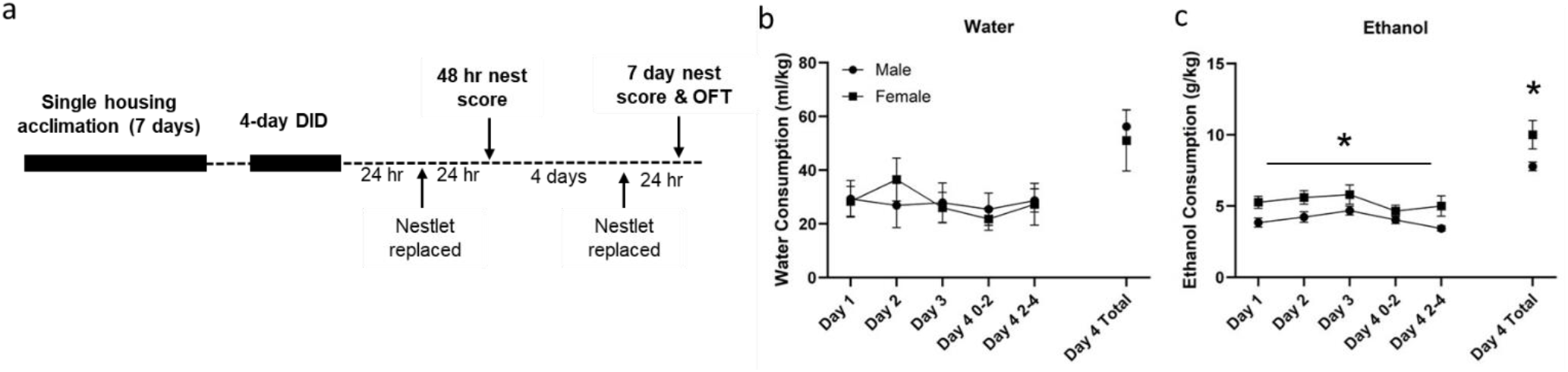
Experiment timeline (A), average ml/kg water consumption of the control animals (B), and average daily g/kg ethanol consumption during the 4-day DID procedure (C). During the drinking procedure, mice had their water bottles switched to sipper tubes containing ethanol or water for 2 hours from days 1 to 3, and for 4 hours on day 4.

**Figure 2.**
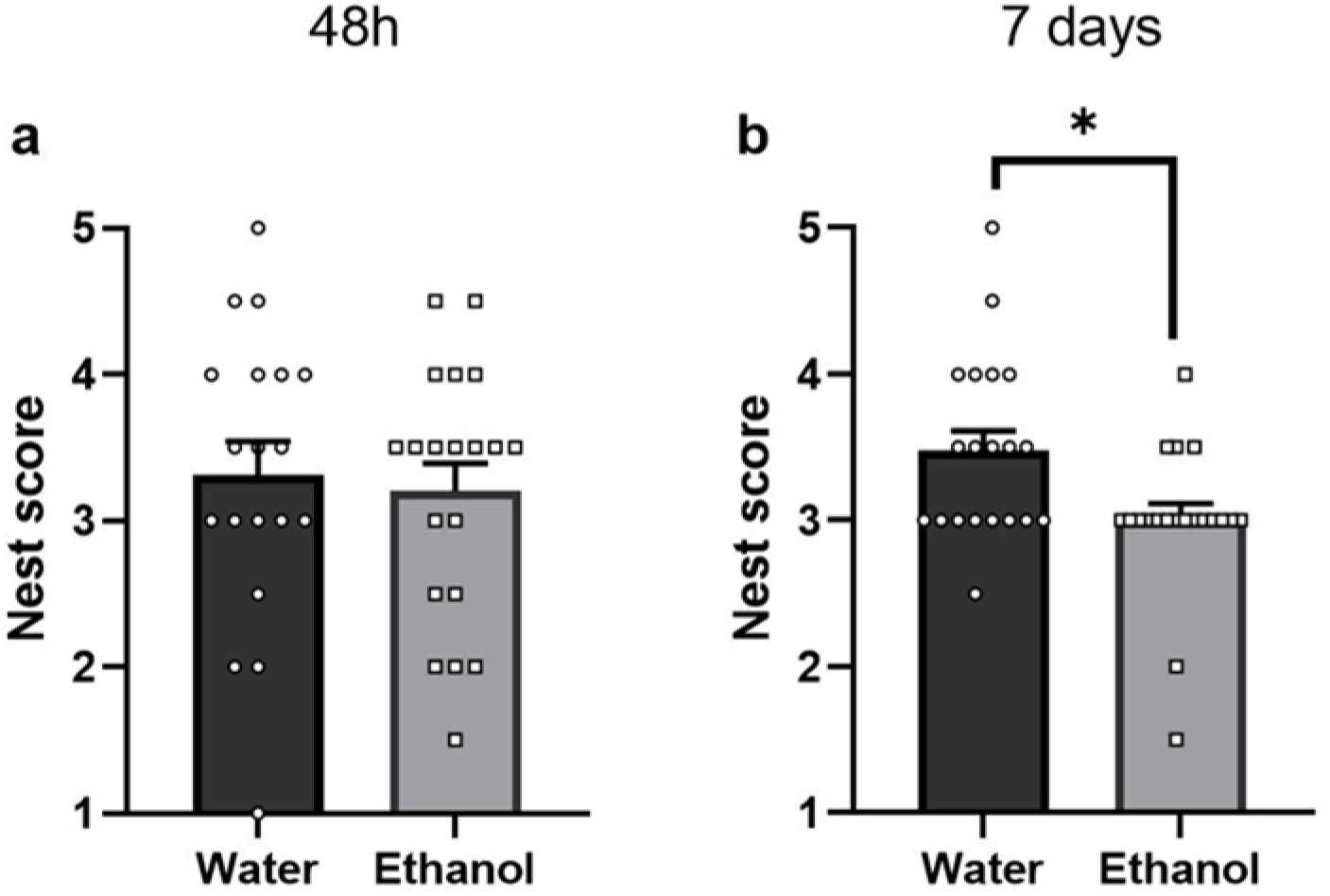
Nest scores at 48h (A) and 7 days (B) after the final DID session. Nests were scored on a scale from 1 to 5, with 5 being the highest in quality. Data are shown as means ± SEM and are collapsed on sex. * indicates *p* < 0.05. N= 10/sex/treatment group.

### Open Field Test

To assess whether deficits in nest building were potentially due to post drinking-associated changes in locomotor activity and anxiety-like behavior, an open field test was performed 7 days after the end of DID (immediately following scoring of nest quality). Figure 3 shows total distance traveled in 15 min (Fig. 3A) and the percent of total time spent in the center of the arena (Fig. 3B). For both variables analyzed, there were no main effects of treatment group and no significant treatment x sex interactions [*F*(1,36) ≤2.64, *p*≥0.113 for all], though there was a significant main effect of sex on center time [males>females; *F*(1,36) =8.58, *p*=0.006].

**Figure 3.**
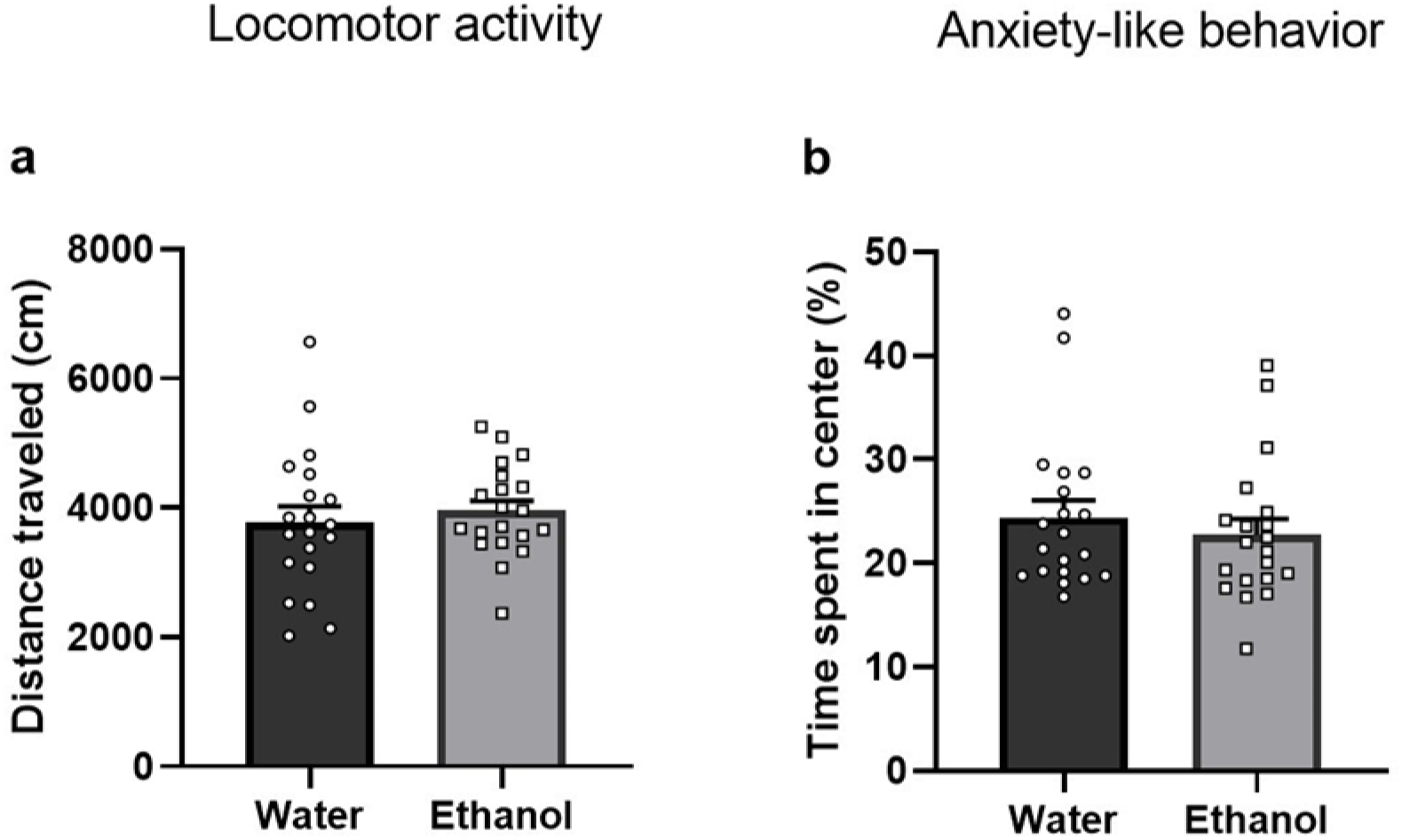
Open field test of locomotor activity and anxiety-like behavior 7 days after the final DID session. Total distance traveled in 15 min (A) and the percentage of total time spent in center (B) are shown. Data are shown as mean ± SEM and are collapsed on sex. N=10/sex/treatment group.

## Discussion

Ethanol withdrawal is a complex state that includes different phenotypes that occur at different timepoints during abstinence. Acute withdrawal typically occurs during the first 48-72 hours after cessation of ethanol in humans (24-48 hours in rodents) and is usually defined by symptoms related to nervous system hyperactivity. In contrast, early abstinence (3-6 weeks in humans, 1-2 weeks in rodents) symptoms may include anxiety-like behaviors and disturbances in sleep with the absence of physical symptoms. Finally, protracted abstinence can last for more than a month and is characterized by the prevalence of negative affective changes^3,16,17^. For all phases of withdrawal, specific symptoms may vary depending on the duration, pattern, and intensity of prior ethanol exposure.

In this study, we found that a non-dependence inducing 4-day DID procedure was sufficient to cause a significant impairment in the nest quality of ethanol-drinking C57BL/6J mice. This impairment was not seen 48 hours after the last drinking session (early post-drinking period), but was significantly different from water-drinking control animals at 7 days after the end of DID (late post-drinking period). This is some of the first evidence for ethanol drinking-associated impairment of nest building during early abstinence following voluntary binge-like drinking. The absence of a drinking group difference during this early post-drinking period is consistent with a previous study which found no differences in nest scores at 24 hours after the last day of ethanol exposure in mice that received a two-bottle choice drinking paradigm^18^.

Interestingly, several other studies have found deficits in nest building during this early post-ethanol time period following acute injections or vapor inhalation, though these studies also used different mouse strains^4,11^. This suggests that route of ethanol administration and genetic background may impact the timing of post-ethanol and withdrawal-related effects on nest building deficits. Another factor that could potentially impact nest quality is sex. Both male and female mice produce nests^11^, though there is some evidence of sex differences in nest scores at baseline ^9^, after ethanol injections^4^, and in aging mice^8^. These differences are not consistent across studies and likely are affected by genetic background^19^. For example, naïve male mice from a heterogeneous stock had higher nest scores than females^11^, whereas several studies in C57BL/6 mice showed no sex differences in nest quality in young adult mice^8,9^, but there is evidence of poorer nest quality in aged males (25 months of age)^8^. Our current findings appear to be consistent with these reports of no sex differences in young adult C57BL/6 mice, but we cannot rule out the possibility of sex x ethanol interactions that may emerge later in the lifespan.

We found no significant effect of drinking group (ethanol vs. water) on locomotor activity and anxiety-like behavior in the open field test when tested 7 days after DID. This suggests that the group difference in nest building performance at this timepoint was likely not due to underlying locomotor deficits or alterations in anxiety level. While locomotor sedation has been commonly reported during early post-ethanol timepoints in mice and could potentially explain reduced nest building activity^3,20^, these locomotor effects are typically gone by the 7-day timepoint during which we observed impaired nest building. This suggests that the time course of locomotor deficits and nest building deficits during the post-drinking period in our experiment shared minimal overlap. Alternatively, locomotor impairment could potentially drive the acute withdrawal differences in nest building deficits that have been previously reported following ethanol injection or vapor exposure, but our drinking paradigm may have been insufficient to produce these locomotor effects. Future experiments that test locomotor activity during the early post-drinking period following DID may help to resolve this distinction.

Similarly, examining nest building behavior at later timepoints following acute ethanol injection of vapor inhalation would be useful to determine whether the 7-day post-ethanol nest building deficit we observed here is conserved across different ethanol exposure paradigms. It may be that nest building deficits following ethanol treatment are due to several different underlying causes, with early post-ethanol deficits reflecting locomotor or thermoregulatory changes during this time period, whereas later post-ethanol deficits might be more closely related to cognitive impairments or negative affective states that predominate during this phase of early abstinence.

There are some potential limitations of the present study that should be noted. First, nest building behavior was only evaluated at two timepoints that were relatively far apart (48 hours and 7 days after the last drinking session). Our primary goal was to determine the effect of binge-like drinking on nest building behavior during the initial post-drinking period vs. a later timepoint when early “hangover”-type symptoms are expected to have resolved. We therefore were not able to determine when the impaired nest building first occurs, or for how long this deficit may persist. Testing a more complete time course of nest building behavior throughout the early post-drinking period and beyond the 7-day timepoint will be necessary to identify more subtle changes in this trait throughout the initial phases of ethanol abstinence. It will also be important to evaluate whether subsequent drinking challenges produce the same effects as an initial DID exposure. Additionally, without BECs we cannot say definitively that the mice reached binge levels of intake. Although ethanol consumption was high and was consistent with levels that have been shown to produce intoxication in previous studies^13-15^, future experiments including a blood sample for BEC determination after the final DID session would be beneficial. We could then also determine whether BEC is correlated with nest building scores, even in the absence of a correlation between total consumption and nest quality. Finally, although we did not see evidence of treatment differences in anxiety-like behavior in the open field test, it is possible that other assays (e.g. elevated zero maze, light-dark box, marble burying) might yield different results.

In summary, our findings provide some of the first evidence for lasting effects of voluntary binge-like ethanol drinking on nest building behavior in mice. These results add to the previous literature showing nest building deficits can serve as an index of acute ethanol withdrawal severity following experimenter-administered ethanol and support the use of this test to evaluate behavioral changes during early abstinence. Because of its ease of use, minimal invasiveness, and relevance for naturalistic mouse behavior, nest building should be considered as a potential phenotype of interest in future studies of ethanol and other drug withdrawal. Increasing the number and type of tasks used to assess post-drinking behavioral changes will ultimately lead to a better understanding of the neurobiology of ethanol withdrawal.

## Data availability

The data collected and analyzed in this study are available on reasonable request from the corresponding author.

## Acknowledgements

This work was supported by a grant from the National Institute on Alcohol Abuse and Alcoholism to AMB-L (R00AA027835).

## Author contributions

ALS-B was responsible for data acquisition, data analysis, and writing the original manuscript draft. AMB-L was responsible for experimental conception and design, data analysis, review and editing of the manuscript, and funding acquisition.

